# InferPloidy: Fast and accurate ploidy inference enables inter-tumoral biomarker discovery in single-cell RNA-seq datasets

**DOI:** 10.1101/2025.03.13.643178

**Authors:** Dawon Hong, Wonjung Sung, Jaeyoung Chae, Jucheol Moon, Jong-chan Lee, Sunjoo Jeong, Seokhyun Yoon

## Abstract

**Background:** Accurate inference of copy number variation (CNV) and ploidy from single-cell RNA-seq data is essential for resolving tumor heterogeneity and identifying malignant cells, yet existing tools such as CopyKat and SCEVAN are limited by long runtimes and reduced accuracy in large or heterogeneous datasets.

**Results:** Here, we present InferPloidy, a high-speed and robust ploidy inference method built on InferCNV that combines graph-based cell-grouping with iterative Gaussian mixture modeling. Across multiple cancer types—breast cancer, non-small cell lung cancer, pancreatic ductal adenocarcinoma, and colorectal cancer—InferPloidy achieved up to two orders of magnitude faster runtimes than existing tools, while maintaining superior classification accuracy. This accurate separation of aneuploid tumor cells enabled the discovery of subtype-specific therapeutic targets, including *ERBB2*, *ESR1*, *EGFR*, and *MET*, as well as recurrent surfaceome markers such as *CD82*, *F11R*, *SLC2A1*, *TM9SF2*, *CXADR*, and *PLPP2*, several of which have preclinical or clinical relevance.

**Conclusion:** These results establish InferPloidy as a scalable platform for CNV-guided tumor cell identification and surfaceome-based biomarker discovery, offering broad utility for precision oncology and translational research.

## Background

Single-cell RNA sequencing (scRNA-seq)^1,2,3^ is a powerful technique for dissecting cancer tissues and the tumor microenvironment ^4^—a complex ecosystem composed of tumor cells, immune cells, and stromal subsets^5,6,7,8^. A major challenge in scRNA-seq-based TME studies is understanding intra and inter tumoral heterogeneity^9^ to get critical insights into tumor evolution and progression^10, 11,12^.

A crucial step in scRNA-seq data analysis for studying TME and TH is the precise classification of tumor cells, not only from normal cells (including immune and stromal cells) but also from normal tumor-originating cells, e.g., epithelial or ductal cells. Accurately distinguishing between tumor and normal cells enables researchers to explore tumor heterogeneity, investigate carcinogenesis, and study tumor-normal cell interactions within the TME^13,14,15^. Furthermore, scRNA-seq analysis of cancer tissues has been instrumental in uncovering mechanisms of drug resistance, identifying novel biomarkers, and exploring therapeutic strategies^16,17,18^.

Several computational tools have been developed for estimation of copy number variation (CNV) and tumor cell classification at the single-cell level, including CopyKat^19^ and SCEVAN^20^. Other related tools, such as InferCNV, developed in a series of studies^21–23^ (https://github.com/broadinstitute/infercnv/wiki and https://github.com/icbi-lab/infercnvpy), and sciCNV^24^, focus solely on CNV estimation. Both CopyKat and SCEVAN provide CNV estimates along with ploidy classification, distinguishing cells as either diploid or aneuploid^25, 26,27^. Accurate CNV estimation allows researchers to study intra-tumoral heterogeneity and tumor evolution by identifying clonal substructures and their phylogeny^10, 11,12^.

However, a major limitation of these tools is their computational inefficiency. While SCEVAN is somewhat faster than CopyKat, both require hours to process moderate-sized datasets (tens of thousands of cells) and days for large datasets (hundreds of thousands of cells). This runtime is significantly longer compared to other scRNA-seq analysis tools used for cell type annotation, cell-cell interaction inference, and differentially expressed gene (DEG) analysis, making CNV-based tumor classification a computational bottleneck and limiting its integration into comprehensive single-cell analysis pipelines.

To address these challenges and enable rapid analysis of large scRNA-seq datasets, we introduce InferPloidy, a high-speed and highly accurate ploidy inference tool that runs on top of InferCNV. InferPloidy facilitates the identification of tumor heterogeneity (TH) across individuals and cancer types, enabling pan-cancer heterogeneity studies and supporting cancer subtyping for biomarker discovery and drug target identification.

### Implementation

The detailed processing steps of InferPloidy are as follows

### Cell type annotation and CNV estimation

First, cell type annotation was performed using HiCAT^28^, which assigns cell identities across three hierarchical levels: major, minor, and subset. Major cell types—including T cells, B cells, myeloid cells, and fibroblasts—were selected and used as normal reference cells. These reference cells were then supplied to InferCNV^21^ to infer copy number variation (CNV) profiles for all cells, using the hg38 gene annotation (GTF format). When no reference cells are available, InferCNV is run in reference-free mode.

### Clustering and building cluster adjacency graph

Dimension reduction was performed on the CNV profiles using PCA with 15 components. Then, using PCA-transformed CNV patterns, k-neighbors graph, *G∼*, was built to perform Louvain clustering^29^. In *G∼*, each node corresponds to a cell and the edge, *e*_*ij*_, represents connection between two cells (*i*, *j*), which is 1 if *j* is one of the *k* nearest neighbors of *i* and 0 otherwise. Python packages, scikit-learn and scikit-network, were used. With the cluster labels, we next build the cluster adjacency graph, *G*, where a node corresponds to a cluster and an edge to connectivity between them. The connectivity, *c*_*ij*_ between a pair of clusters, (*i*, *j*), is defined as follows:

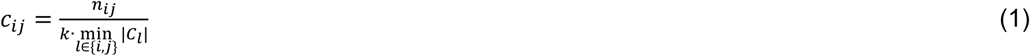

where |*C*_*l*_| is the size (number of cells) of cluster *l* and *n*_*ij*_ is the number of edges in *G∼* connecting cluster *i* and *j*. If a cell in cluster *i* has a nearest neighbor belonging to cluster *j*, it is counted in *n*_*ij*_. With the number *k* in the denominator, *c*_*ij*_ satisfies 0 ≤ *c*_*ij*_ ≤ 1. The connectivity in (1) represents how close the cluster *i* and *j* are to each other.

### Grouping clusters into three: Normal reference, reachable, and unreachable

First, normal reference clusters are defined as those containing at least 25% of normal reference cells. Then, the following successive procedure is used to divide the rest of clusters into the reachable and the unreachable clusters from normal reference clusters. Let *N* be the set of normal reference clusters and *R* be the rest. The clusters in *R* are ordered by choosing cluster one-by-one as follows.

A. Initialize and empty set *R*^′^ = ∅
B. Select cluster *j*^∗^ ∈ *R* such that 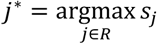 with *s*_*j*_ = ∑_*i*∈*N*∪*R*′_ *c*_*ij*_,
C. Store the cluster index *j*^∗^ and its connectivity score *s*_*j*_∗, and move *j*^∗^ from *R* to *R*′
D. do step B and C successively until no cluster is left in *R*.
E. By inspecting the ordered set of connectivity scores {*s*_*m*_: *m* = 1,2, …, |*R*|} from the first, find the first cluster *m*^∗^ of which *s*_*m*_∗ < *c*_*th*_ (some connectivity threshold)
F. Reachable clusters set is defined as *U* = {*j*_*m*_: *m* = 1,2, …, *m*^∗^ − 1} while unreachable as *T* = {*j*_*m*_: *m* = *m*^∗^, … ., |*R*|}

Note: If the summation in step B is replaced to maximization, i.e., 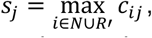 the problem becomes equivalent to ordering graph nodes by their maximum path cost to any one in *N*, where a path cost is defined as the minimum connectivity along the path. In this case, shortest path search procedure, such as Bellman-Ford algorithm, can be applied. However, we used the summation-based definition to mitigate variability due to stochastic clustering behavior, as summation offers more stability across repeated runs and minor dataset perturbations.

If no normal reference cells are present in the dataset, an alternative approach is applied. The largest, tightly connected clusters are first identified and used as pseudo-reference clusters. Then, the same procedure (steps A–D) is applied to classify remaining clusters into reachable and unreachable groups.

### CNV profiling of normal and tumor cells using Gaussian mixture model

Next, our objective is to classify cells in the reachable (U) and unreachable (T) clusters as either normal or tumoral. This classification is based on two key assumptions:

(1) cells in the original reference clusters *N* are reliably normal, and ^30^ cells in the unreachable set *T* are predominantly tumoral, although not exclusively.

Based on this assumption, we model the CNV distribution of normal and tumor cells separately using a Gaussian Mixture Model (GMM). For each genomic position *m*, the probability density function of CNV values for group *X* (where *X* ∈ {*N*, *T*}) is defined as:

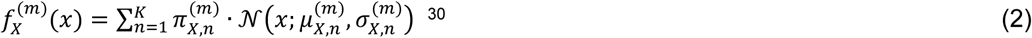

where *K* is the number of Gaussian components, *N*(*x*; *μ, σ*) denotes a normal distribution of mean *μ* and standard deviation *σ*, and 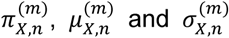 are the mixing weight, mean and standard deviation of the *n*^th^ component for genomic position *m* in group *X*.

Model parameters for both *N* (normal) and *T* groups are estimated using the Expectation-Maximization (EM) algorithm. In our implementation, we used the ‘GaussianMixture’ class from the *scikit-learn* Python package to fit the model and obtain the parameters for each genomic position.

### Decision metric

Given the Gaussian mixture model parameters for the tumor (*T*) and normal (*N*) cell populations, we defined a decision metric to distinguish tumor-like from normal-like CNV profiles. For each cell, the log-likelihood ratio across genomic positions is computed as:

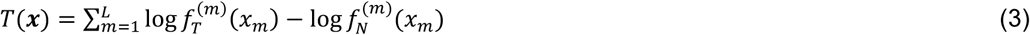

where *L* denotes the number of genomic positions at which CNV’s were evaluated, the vector *x* = [*x*_1_, *x*_2_, …, *x*_*m*_, …, *x*_*L*_] represents the CNV values of a single cell, where *x*_*m*_ corresponds to the CNV at the *m*^th^ genomic position. In our implementation, *L* was set to 3, with each position ideally capturing one of three canonical CNV states: gain, loss, and neutral.

The use of a Gaussian mixture model, rather than a single-component model, is motivated by the inherent heterogeneity in tumor CNV profiles. Multiple subpopulations within a tumor may exhibit distinct CNV states even at the same genomic locus. This rationale also applies to normal cells, as diverse normal cell types—such as immune cells and epithelial cells—can display unique CNV patterns due to lineage-specific gene expression.

It is important to note that the model parameters were initially estimated using cells from the reference-normal (*N*) and presumed-tumor (*T*) clusters. Therefore, it is not guaranteed that these models will generalize well to the *reachable* cluster group (*U*), which was identified as distinct from both *N* and *T* in the initial clustering. Cells in *U* may include other normal cell types (e.g., endothelial or epithelial cells not represented in *N*), as well as tumor cells that exhibit CNV patterns differing from those in *T*. To address this uncertainty, we implemented an iterative detection and re-estimation strategy, rather than making direct classifications based solely on the initial models.

### A cautious approach to make final decision: iterative refinement

Let *N*∼, *T*∼, and *U*∼ denote the sets of cells of currently assigned as normal, tumor, and undecided, respectively. Initially, they are defined by the cells belonging to the clusters in *N*, *T*, and *U*, respectively. The Gaussian mixture model parameters for the normal and tumor groups, denoted as *p_N_* and *p_T_* include all 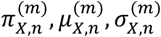 for each component *n* and genomic position *m* (as defined in Equation 2). These are updated iteratively through the following procedure:

A. **Cell set assignment**: Based on the current cluster groups *N*, *T*, and *U*, update the cell-level groupings *N*∼, *T*∼, and *U*∼.
B. **Model parameter estimation**: Recalculate *p*_*N*_ and *p*_*T*_ using CNV profiles from cells in *N*∼ and *T*∼, respectively.
C. **Decision metric computation**: For each cell *j*, compute the decision metric *t*_*j*_ = *T*(*x*_*j*_), as defined in Equation (3), using the updated model parameters.
D. **Threshold calculation**: Compute the mean and the standard deviation of *t*_*j*_ for the normal cells in *N*, (*μ*_*N*_, *σ*_*N*_), and those for tumor cells in *T*∼, (*μ*_*T*_, *σ*_*T*_). Then, define two decision thresholds: *th*_*lower*_ = *μ*_*N*_ + α · *σ*_*N*_ and *th*_*upper*_ = *μ*_*T*_ − α · *σ*_*T*_, where 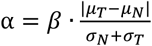 with *β* being a hyperparameter between 0 and 1 that controls the decision margin.
E. **Cell reclassification**: Given the upper and lower threshold, the cells with *t*_*j*_ < *th*_*lower*_ are decided as normal, those with *t*_*j*_ > *th*_*upper*_ as tumor, and others are set ‘undecided’. These decisions are applied only to those cells in *T*∼ and *U*∼.
F. **Update cell-level grouping**: Update *N*∼, *T*∼, and *U*∼ as follows:

a. *N*∼ ← *N*∼ ⋃ {*j*: *t*_*j*_ < *th*_*lower*_ ∀*j* ∈ *T*∼ ∪ *U*∼},
b. *T*∼ ← {*j*: *t*_*j*_ > *th*_*upper*_ ∀*j* ∈ *T*∼ ∪ *U*∼}
c. *U*∼ ← {*j*: *th*_*lower*_ ≤ *t*_*j*_ ≤ *th*_*upper*_ ∀*j* ∈ *T*∼ ∪ *U*∼}
G. **Update cluster-level grouping**: Using the newly updated *N*∼ and the cluster labels previously assigned to each cell, redefine the cluster-level groups *N*, *T*, and *U* based on the following:

i. Clusters with ≥ 25% of their cells in *N*∼ are assigned to *N*
ii. Reachable clusters from (the updated) *N* are assigned to *U*
iii. Unreachable clusters are assigned to *T*. (In this step, only cluster grouping is changed without altering the cell’s cluster membership and the reachability between clusters.)
H. **Repeat**: The steps A to G are repeated until the cluster assignment *N*, *T*, and *U* no longer changes or a predefined maximum number of iterations is reached.

**Note:** Since the number of normal cells in *N*∼ is non-deceasing across iteration, the process can be ended up with all cells decided as normal if the initial grouping is overly biased toward normal cells. Thus, careful initialization of *N*∼, *T*∼, and *U*∼ is essential to ensure biologically meaningful classification.

### Consideration in hyper parameter setting

The initial classification of cells into *N*∼, *T*∼, and *U*∼ is contingent upon the initial clustering of cells into *N*, *T*, and *U*, which depends on the connectivity threshold *c*_*th*_, used in the graph-based cluster partitioning. Careful selection of *c*_*th*_ is therefore critical for accurate designation of normal reference clusters. Based on empirical evaluation across multiple datasets, we found *c*_*th*_ = 0.18 to perform well in most cases. For datasets with stronger inter-cluster connectivity’s, particularly where improved true negative rates (i.e., correct identification of normal cells) are desired, a higher threshold may be beneficial. In this study, the threshold was adaptively set using inter-cluster connectivity statistics from clusters in *N*, i.e., *c*_*th*_ = max (0.18, *μ*_*C*_ − 1.5*σ*_*C*_) with *μ*_*C*_ and *σ*_*C*_ denote the mean and the standard deviation of the connectivity values among clusters in *N*.

Another important consideration is the inherent stochasticity of clustering algorithms. Even with fixed hyperparameters, different clustering seeds can lead to varied outcomes due to random initialization. While fixing the seed may yield reproducible results, it may not capture the representative outcome from a distribution of possible clusterings. This variability is not unique to our method, but a general challenge in unsupervised learning. To mitigate this, we performed the inference procedure across multiple clustering runs and applied majority voting at the cell level to determine final classifications. This ensemble approach improves robustness and yields a more balanced classification between normal and tumor cells, reducing the risk of biased assignments stemming from any single clustering outcome.

Lastly, the hyperparameter *β*, which determines the decision margin for classification, must also be set judiciously. A large *β* yields conservative classification with many undecided cells, whereas a small *β* risks imbalanced outcomes by overly favoring either normal or tumor designation. In our framework, *β* was initialized to 0.4 and reduced by 0.1 in each iteration of the refinement loop (steps A to H), until it reached a minimum value of 0.2.

### Sample diversity index

To quantify inter-sample heterogeneity within each cluster, we computed a Sample Diversity Index (SDI) using Shannon’s entropy, defined as:

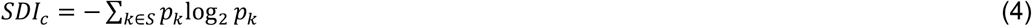

where *S* is the set of sample indices and *p*_*k*_ the proportion of cells originating from the *k*^th^ sample within cluster *c*. This metric is equivalent to the Shannon diversity index commonly used in microbiome studies. By its mathematical property, the SDI ranges between 0 and log_2_|*S*|, where |*S*| is the total number of samples. A value of 0 indicates that the cluster is composed entirely of cells from a single sample, while values approaching log_2_|*S*| suggest a well-mixed cluster containing cells from many different samples.

### Running CopyKat and SCEVAN

CopyKat and SCEVAN were executed using their respective default hyperparameters, with the exception of specifying the normal reference cells. For consistency across tools—including InferCNV and InferPloidy—T cells, B cells, myeloid cells, and fibroblasts were designated as normal reference populations.

### Assessment of marker scores

For each dataset, we first identified the tumor-originating cell type—for example, epithelial cells in breast cancer, NSCLC, and CRC, and ductal cells in PDAC. Cells of the tumor-originating type were then classified into three groups: (i) normal (or adjacent normal) cells from normal or adjacent normal samples, (ii) diploid cells from tumor samples, and (iii) aneuploid cells from tumor samples. Aneuploid cells were further subdivided based on clinical or molecular features; for instance, ER⁺, HER2⁺, and triple-negative breast cancer (TNBC) subtypes in the breast cancer dataset (GSE161529), and ‘early tumor’ and ‘advanced tumor’ stages in the NSCLC dataset (GSE131907).

Based on these groupings, differential gene expression (DEG) analysis was performed to calculate the proportion of cells expressing each gene in the target group (denoted as pct_nz_test) and in the reference group (all other groups, denoted as pct_nz_reference). The marker score for each gene was defined as follows:

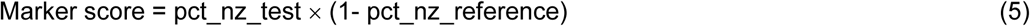

Genes were ranked according to their marker scores, and the top M genes were selected from among those encoding cell surface proteins (i.e., surfaceome genes) for downstream analysis.

### Identifying the genomic spots with significant CNV gain

From the tumor-originating cell types (ductal cells for PDAC and epithelial cells for other cancers), we first collected all diploid cells to calculate the mean *μ*_*d*_(*p*) and standard deviation *σ*_*d*_(*p*) for each genomic spot *p*. For each sample *s*, we then collected aneuploid cells to calculate their mean *μ*_*a*_(*s*, *p*) and standard deviation *σ*_*d*_(*s*, *p*). Instead of using the overall mean of aneuploid cells in sample *s*, we fitted a Gaussian mixture model with a diagonal covariance constraint and selected the largest component mean as *μ*_*a*_(*s*, *p*) and its corresponding standard deviation as *σ*_*a*_(*s*, *p*). This approach accounts for the fact that, unlike normal cells, tumor cells are highly heterogeneous, and not all cells within a sample necessarily share similar CNV patterns. The significance score was defined as:

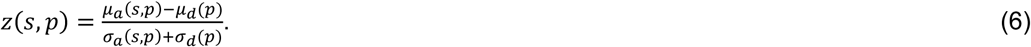

Genomic regions with significant CNV gain were identified by selecting spots with *z*(*s*, *p*) ≥ 0.5. Finally, we retained positions *p* that met this threshold in at least *N* samples (*N* = 4 in this study) and calculated *z*(*p*) as the mean of *z*(*s*, *p*) across those samples. (**Fig. 9**)

## Results

### Overview of InferPloidy

InferPloidy utilizes pre-computed CNV estimates and a predefined set of normal reference cells. To achieve this, we first use HiCAT^28^ for cell-type annotation and then collect T cells, B cells, myeloid cells, and fibroblasts to use as normal reference cells for InferCNV (**Fig. 1a, b**). The core idea is to use CNV estimates and predefined normal reference cells to initially classify cells into three groups: normal, tumoral, and uncertain (others). Using the first two groups, CNV profiles for normal and tumor cells are computed, allowing an initial classification not only for the uncertain cells but also for tumoral cells.

**Figure 1.**
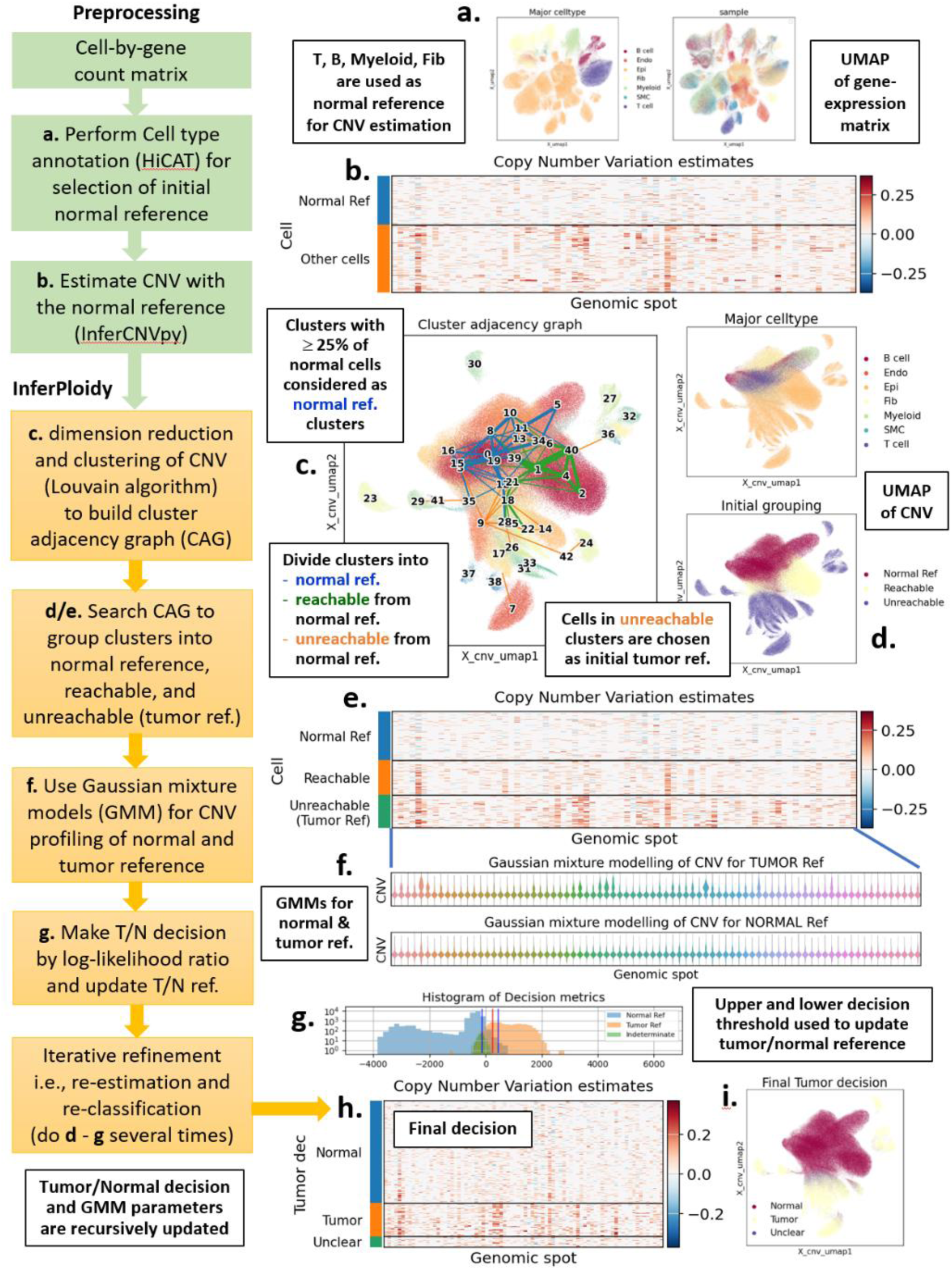
A brief description of InferPloidy processing steps. **a**, **b:** It starts with CNV estimation using InferCNV, for which HiCAT was used for annotation of cell types to collect normal reference cells. **C:** With the CNV estimates, dimension reduction using PCA and Louvain clustering are performed to find cluster adjacency graph. **d**, **e:** By searching the graph, clusters are divided into three groups, i.e., normal reference clusters, clusters reachable from normal reference, and those unreachable, where the cells in the unreachable clusters are initially used as tumor reference cells. **f:** The CNV patterns of normal and tumor reference cells are modelled by using Gaussian mixture model with diagonal covariance constraint. **g:** The initial ploidy decision is made by the log-likelihood ratio test. Step **d** to **g** is repeated until no changes in ploidy decision is found or the number of repetitions reaches the predefined number.

Specifically, we first perform dimensionality reduction (using PCA) and clustering on the CNV estimates to construct a cluster adjacency graph (CAG) (**Fig. 1c**). The CAG can be efficiently generated using a graph-based clustering algorithm, such as Louvain clustering^29^. Based on this CAG, clusters are divided into three categories: (i) the normal reference clusters, (ii) the clusters that are “reachable” from the normal reference clusters, and (iii) those “unreachable” from the normal reference clusters (**Fig. 1d**). To this end, the normal reference cluster is first defined as the cluster where the proportion of normal reference cells (T cells, B cells, and myeloid cells) exceeds a certain threshold. With the normal reference clusters, we next assess reachability from the normal reference clusters to classify the remaining clusters into “reachable” and “unreachable”. Clusters unreachable from the normal reference clusters are initially designated as tumor reference clusters while those reachable are set “uncertain”. Using these normal and tumor reference clusters, the CNVs of normal and tumor cells are approximated via a Gaussian Mixture Model (GMM) with a diagonal covariance constraint, where model parameters are estimated using the Expectation-Maximization (EM) algorithm (**Fig. 1e, f**).

Ploidy decisions are then made on the cells in the tumor reference clusters and the uncertain clusters by computing the likelihood ratio between the GMMs of normal and tumor cells (**Fig. 1g, h, i**). Steps d to g are iteratively repeated until either the ploidy classifications remain unchanged or a predefined maximum number of iterations is reached (inner loop for iterative refinement). This entire process, from step c to g (including the inner loop), is also repeated multiple times (outer loop) to obtain a final ploidy classification via majority voting. The outer loop ensures robustness by aggregating results from clustering runs with different random initializations.

A detailed description of the InferPloidy workflow can be found in the Implementation section.

### Qualitative comparison of InferPloidy and other tools

We first compared the ploidy classification of InferPloidy with that of CopyKat and SCEVAN using the lung cancer dataset GSE131907^31^ (**Fig. 2 a-c**). The original dataset consists of 58 samples, including normal lung, tumor lung, lymph node, and brain metastases, totaling 208,000 cells. To reduce the runtime of CopyKat and SCEVAN (but not InferPloidy), we excluded lymph node and brain metastasis samples and randomly selected up to 1,200 cells per sample from the remaining three conditions (distant normal, early tumor, and advanced tumor), analyzing a total of 36,000 cells without splitting them into individual samples. For a fair comparison, all tools used the same set of normal reference cells, including T cells, B cells, myeloid cells, and fibroblasts.

**Figure 2.**
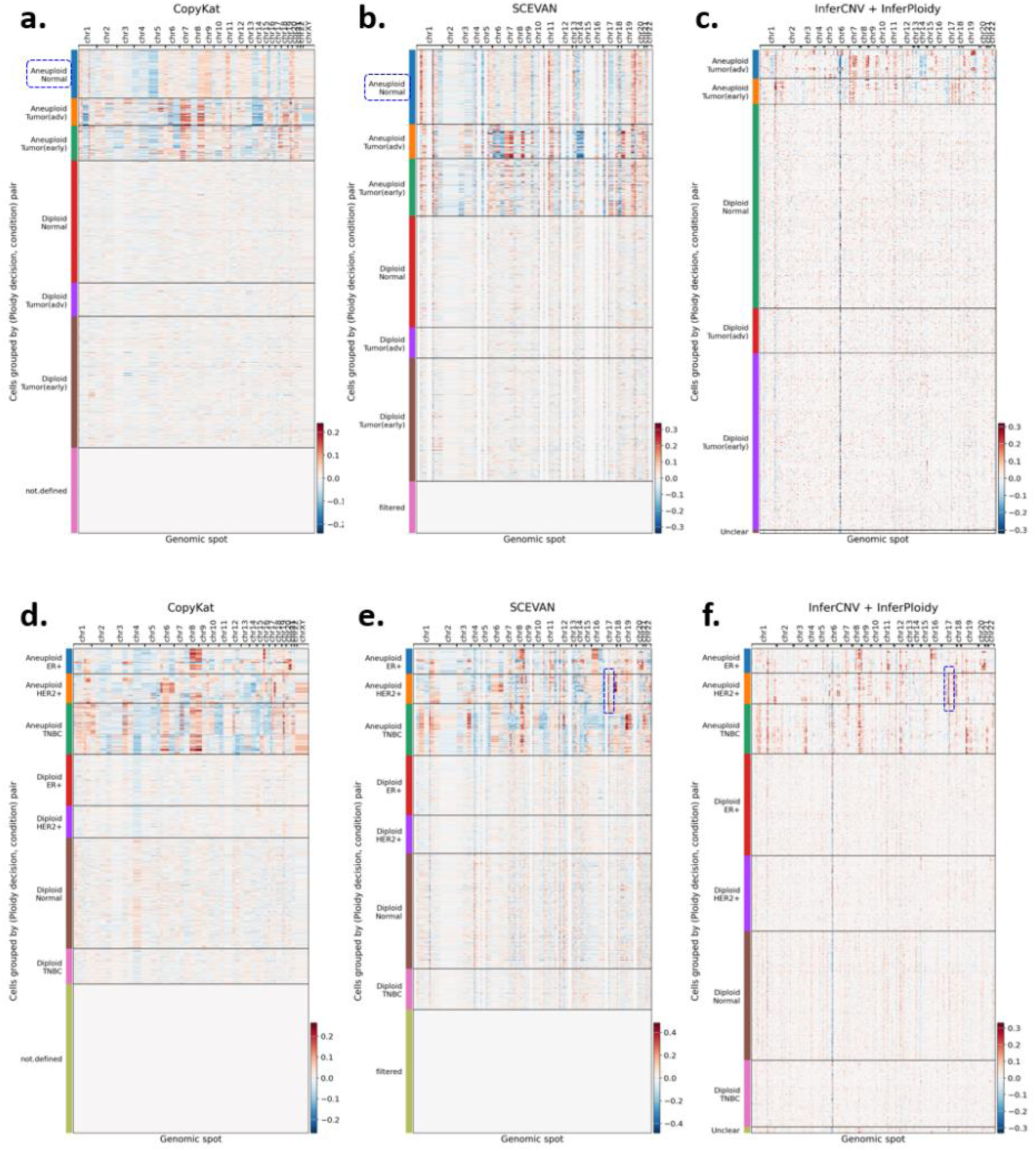
A comparison of CNV estimates and ploidy decisions for lung cancer dataset GSE131907*, which includes three conditions: distant normal, early tumor, and advanced tumor, and for breast cancer dataset GSE176078*, which includes three conditions: ER^+^, HER2^+^, and TNBC (**d-f**). **a,d**: CopyKat, **b,e**: SCEVAN, and **c,f**: InferCNV + InferPloidy. For GSE131907*, many cells were erroneously classified as aneuploid by CopyKat and SCEVAN, whereas no such errors were observed with InferCNV + InferPloidy. In GSE176078*, a copy number gain in the genomic region on chromosome 17, where the ERBB2 gene is located, was identified in the HER2^+^ condition with SCEVAN and InferCNV (blue boxes in **e** and **f**). In both datasets, a considerable portion of cells were classified as indeterminate with CopyKat and SCEVAN.

When classifying cells into aneuploid and diploid groups, we observed that CopyKat and SCEVAN misclassified a non-negligible proportion of cells from the normal condition as aneuploid, which is a clear classification error, as all cells from the distant normal condition should be classified as diploid. In contrast, InferCNV + InferPloidy did not exhibit such errors, demonstrating its superiority in ploidy inference. Another drawback of CNV estimation in CopyKat and SCEVAN was that CNVs for a considerable portion of cells were indeterminate, indicated as either ‘not.defined’ or ‘filtered’ (**Fig. 2 a, b**). This reduces the number of cells usable not only for ploidy inference but also for downstream analyses, such as marker or druggable target discovery. Similar tendencies could also be observed in another dataset, GSE161529 for breast cancer (**Fig. 2 d-f**), where we have collected 36K cells from 26 samples in a similar way to GSE131907.

The UMAP plots provide a detailed visualization of the ploidy classifications from the three tools (**Fig. 3**). Note that the cells in the region where normal reference cells are predominant are likely to be normal, as this region encompasses all clusters containing normal reference cells (**Fig. 3**, normal reference indicator), including T cells, B cells, myeloid cells, and fibroblasts. Within this UMAP region, CopyKat and SCEVAN misclassified many cells as aneuploid, whereas InferPloidy did not. These erroneous classifications in CopyKat and SCEVAN (**Fig. 3a, b**) were mostly myeloid cells from the normal condition that were part of the normal reference set (**Fig. 3d, e, f**).

**Figure 3.**
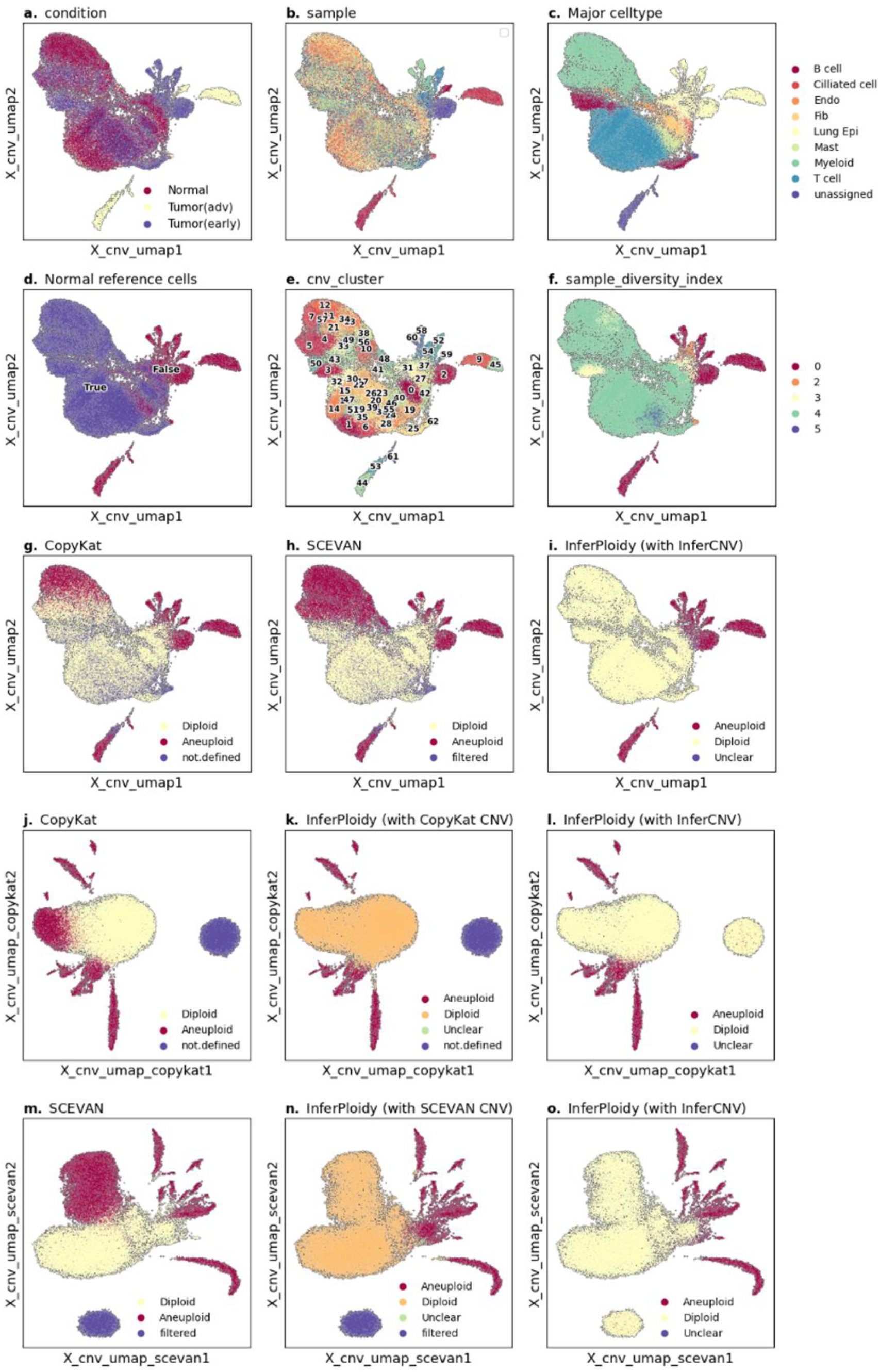
A summary on UMAP projection based on the CNVs obtained by InferCNV, CopyKat (**j-l**), and SCEVAN (**m-o**) for GSE131907*. InferPloidy (with SCEVAN) and InferPloidy (with CopyKat) are the results of InferPloidy using the CNVs obtained by SCEVAN and CopyKat, respectively.

This result may be attributed to either the superior CNV estimation of InferCNV or the superior ploidy inference of InferPloidy. To isolate the contribution of InferPloidy, we applied InferPloidy using the CNV estimates generated by CopyKat and SCEVAN (**Fig. 3j-o**). Even in these cases, InferPloidy successfully classified normal cells from the normal condition as diploid, confirming its superiority in ploidy classification.

Notably, the UMAP region corresponding to normal reference cells aligns with clusters exhibiting high sample diversity index values (**Fig. 3 f and i**). Due to the high heterogeneity of tumor cells, their CNV patterns are often unique to individual samples, resulting in a low sample diversity index—i.e., clusters are predominantly occupied by aneuploid cells from a limited number of samples. In contrast, normal cells exhibit cell-type-specific gene expression patterns, and their corresponding CNV estimates remain relatively unchanged across samples, leading to a high sample diversity index—i.e., normal cells from different samples cluster together. Under InferPloidy’s classification, we observe that diploid and aneuploid cells can be roughly separated based on their sample diversity index values.

### Quantitative comparison

We utilized four datasets with available cancer annotations: GSE176078^32^, GSE103322^33^, GSE72056^30^, and GSE131907^31^, where, in GSE131907, cancer cell annotations are provided only for advanced tumors; therefore, we excluded samples corresponding to early-stage tumors from our analysis. To show the superiority of InferPloidy, we also performed it with the CNV estimates from CopyKat and SCEVAN.

The results indicated that all three tools performed similarly for GSE103322 and GSE72056. However, for GSE176078 and GSE131907, InferCNV + InferPloidy outperformed CopyKat and SCEVAN, achieving an accuracy of 97.7% and 99.5%, respectively and F1-scores of 98.1% and 99.6, respectively. In contrast, CopyKat and SCEVAN yielded much lower accuracy (< 76% and < 65%, respectively) and F1-scores (< 85% and < 74%, respectively) (**Fig. 4a, b**).

**Figure 4.**
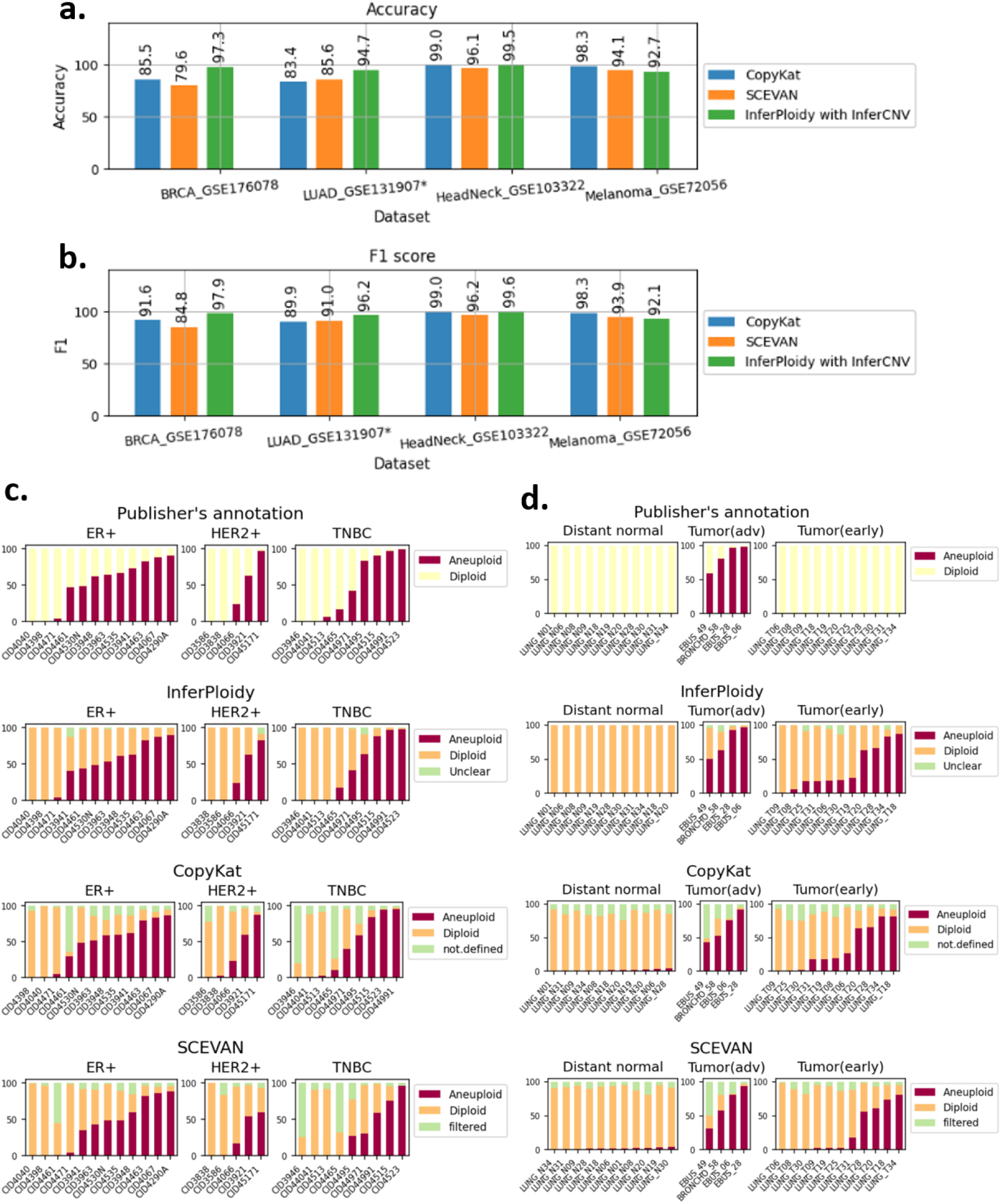
A comparison of accuracies and F1-scores of the ploidy decisions for the datasets with publisher’s ploidy annotations (**a, b**) and a summary of the ploidy decisions per sample for GSE176078* and GSE131907*. The normal reference cells were excluded in the performance evaluation and the summary of per-sample ploidy decisions.

When comparing the proportion of aneuploid, diploid, and ‘undecided’ cells in each sample, CopyKat and SCEVAN exhibited a considerable fraction of ’undecided’ cells (categorized as either not.defined or filtered), whereas this proportion was significantly lower with InferCNV + InferPloidy (**Fig. 4c, d**). As noted previously, in GSE131907, CopyKat and SCEVAN made erroneous classifications and yielded ’undecided’ cells in samples from the distant normal condition. In contrast, InferCNV + InferPloidy correctly classified all cells in distant normal samples as diploid (**Fig. 4d**, distant normal).

One possible concern is that InferCNV + InferPloidy might introduce a bias toward classifying cells as diploid. However, this does not appear to be the case, as InferCNV + InferPloidy identified a greater number of aneuploid cells in both datasets compared to the other tools.

### Running time

One of the most important measures of a software tool is its runtime efficiency in generating final results. While runtime may not be a critical factor if it is relatively short (e.g., on the order of minutes), it becomes a significant concern when the execution time is excessively long. For the down-sampled LUAD dataset from GSE131907, which contains 36,400 cells, CopyKat and SCEVAN required 42 hours and 14 hours, respectively. In contrast, InferCNV and InferPloidy completed the analysis in just 7.5 minutes, respectively—a substantial improvement over CopyKat and SCEVAN.

For a much larger dataset, GSE161529, which comprises 69 samples and 428,000 cells, InferCNV and InferPloidy completed the analysis in less than 2 hours, respectively. In contrast, CopyKat and SCEVAN required 3 days 6 hours and 24 hours, respectively, when each sample was processed separately. In this full dataset, CopyKat and SCEVAN were run one sample at a time, as recommended by the CopyKat developers. However, this approach posed a challenge: some normal samples rarely contained normal reference cells, as immune cells are scarce under some conditions. As a result, CNV estimation tools could not leverage CNV estimates of normal reference cells to accurately infer the CNVs. To address this issue, we randomly sampled cells across the dataset and processed them together especially to get best results of CopyKat and SCEVAN. In contrast to CopyKat and SCEVAN, we executed InferCNV and InferPloidy without dividing the dataset into individual samples. The results demonstrated that InferPloidy consistently outperformed CopyKat and SCEVAN in ploidy inference, even for the down-sampled dataset.

### Accurate ploidy classification enhances subtype-specific and pan-cancer therapeutic target discovery

To highlight the applicability of InferPloidy to investigate inter-tumoral heterogeneity, we tried to identify tumor-specific surface markers and potential therapeutic targets from public single-cell RNA-seq datasets across four major cancer types: non-small cell lung cancer (NSCLC), breast cancer (including triple-negative, HER2-positive, and ER-positive subtypes), pancreatic ductal adenocarcinoma (PDAC), and colorectal cancer (CRC). CNV-based classification enabled accurate separation of aneuploid tumor cells from diploid normal-like cells, providing a robust framework for downstream expression-based target discovery.

We focused our analysis on the surfaceome, defined as the set of genes encoding cell surface proteins with therapeutic or diagnostic relevance. Within the tumor cell populations identified by InferPloidy, we performed differential gene expression analysis to compute marker scores (see Assessment of marker scores) and selected the surfaceome genes up to 50 with the minimum score of 0.25 for each cancer type/subtype. (**Fig. 5, Supplementary Table.1a-f**)

**Figure 5.**
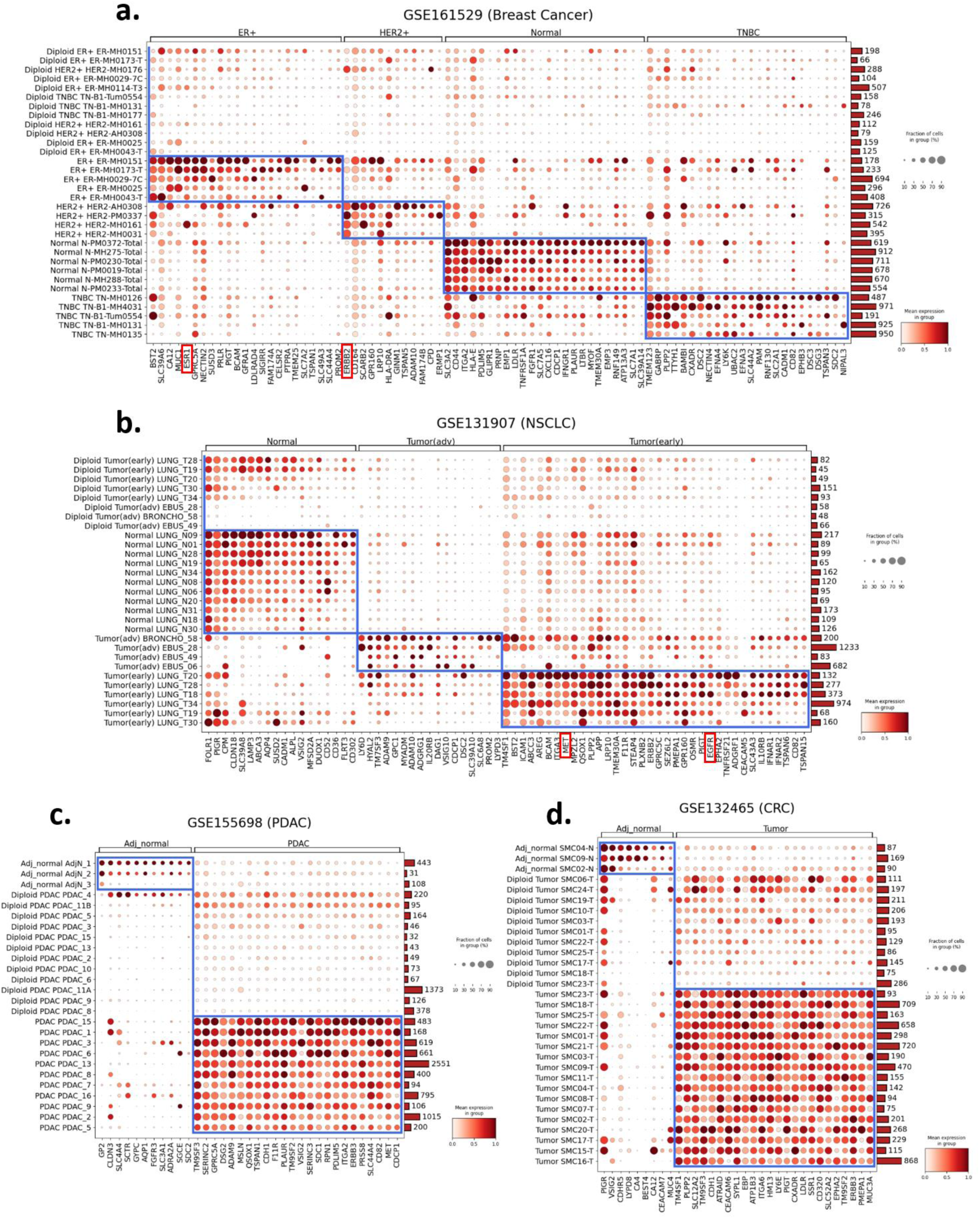
Dot plots showing marker expression patterns across four cancer types: (a) breast cancer, (b) non-small cell lung cancer (NSCLC), (c) pancreatic ductal adenocarcinoma (PDAC), and (d) colorectal cancer (CRC). In each panel, rows represent cell groups from individual samples, stratified into normal (or adjacent normal), diploid tumor, and aneuploid tumor populations. Both diploid and aneuploid cells are derived from tumor samples, with only diploid cells explicitly prefixed as such. In contrast, normal or adjacent normal cells are from normal or adjacent normal samples in which all cells were identified as diploid (by InferPloidy). Columns represent marker genes selected and ranked by their marker scores, as described in the Implementation section. Up to 40 top markers were shown for NSCLC, and up to 25 for the other cancer types. Dot size indicates the fraction of cells in each group expressing the gene, while color intensity reflects the mean expression level. Notably, *ERBB2* and *ESR1* were identified as prominent markers in HER2^+^ and ER^+^ breast cancer, respectively, and *EGFR* and *MET* were highly expressed in NSCLC. These results highlight the utility of InferPloidy in identifying subtype-specific markers and characterizing inter-tumoral heterogeneity across cancer types.

Notably, among the identified genes were several well-established therapeutic targets—*ERBB2* in HER2^+^ and *ESR1* in ER^+^ breast cancer, *EGFR* in NSCLC—which were robustly detected as amplified and overexpressed specifically in aneuploid tumor cells. Including these well know markers, we also could identify other noticeable surface markers that were reported in recent literature as potential therapeutic target. Specifically, in breast cancer, we identified a distinct repertoire of cell surface markers specific to each molecular subtype. For, markers such as *BCAM*^34^, *BST2*^35^, *CA12*^36^, *ESR1*, and *SLC39A6*^37^ were selectively expressed. In HER2^+^ tumors, upregulated markers included *ERBB2*, *ADAM10*^38^, *SCARB2*^39^, and *CD164*^40^. In triple-negative breast cancer (TNBC), surface proteins such as *CD82*^41^, *CXADR* (CAR), *NECTIN4*^42^, and *LY6K*^43^ were identified as highly enriched. Of note, *ENFA3, ENFA4,* and *EPHB3,* related to the Ephrin signaling pathway, were also identified as markers in TNBC subtype (**Fig. 5a**). Eph receptor tyrosine kinases and their ephrin ligands are known to play critical roles in cancer development and progression^44^. *CXADR* (CAR), the coxsackievirus and adenovirus receptor, has also been identified as a therapeutic target, with antibodies (e.g., 6G10A and the human–mouse chimeric ch6G10A) demonstrating tumor growth inhibition in preclinical models via antibody-dependent cellular cytotoxicity (ADCC) and complement-dependent cytotoxicity (CDC)^45^. These findings provide subtype-specific candidates for targeted therapies or diagnostics in breast cancer.

In NSCLC, surface proteins such as *EGFR*, *CEACAM5*^46^, *F11R*(JAM-A)^47^, *MET*^48^, *BCAM*^49^, *TM4SF1*^50^ and were prevalent *TSPAN5/6*^51^ were prevalent among tumor cell clusters (**Fig. 5b**). For PDAC, notable markers include,*CD8252, CDH153, MSLN54, EMP2*55, *F11R*47, *MET48*, and integrins such as (**Fig. 5c**). In CRC, frequently observed surface markers encompass *CEACAM6*^57^, *EPHA2*^58^, *CDH153, ITGA656, SLC12A2* ^59^, *TM4SF1*^50^, *TM9SF2,* and *SDC1*^60^ (**Fig. 5d**). Of note, *SDC1* (Syndecan-1, *CD138*) is a well-characterized cell surface proteoglycan implicated in tumor progression, metastasis, and chemoresistance, and is targeted by investigational antibody-drug conjugates (ADC) therapies (e.g., indatuximab ravtansine in multiple myeloma and solid tumors)^61^.

As documented in the cited references for each marker, all these markers have known oncogenic roles or are under active investigation as therapeutic targets. This observation highlights the precision of InferPloidy in recovering clinically actionable targets from single-cell transcriptomic data. InferPloidy provides a valuable platform for translational cancer research. Its application facilitates the discovery of both known and novel targets, ultimately contributing to the development of more precise and personalized therapeutic strategies.

To further investigate the translational utility of InferPloidy-based tumor cell classification, we compared surface marker gene sets across four cancer types—TNBC, NSCLC, PDAC, and CRC—using the top 40 surfaceome genes identified per cancer type. Through this comparison, we identified a subset of genes recurrently upregulated in tumor epithelial cells across multiple cancers (**Fig. 6**), which may serve as pan-cancer therapeutic targets or biomarkers.

**Figure 6.**
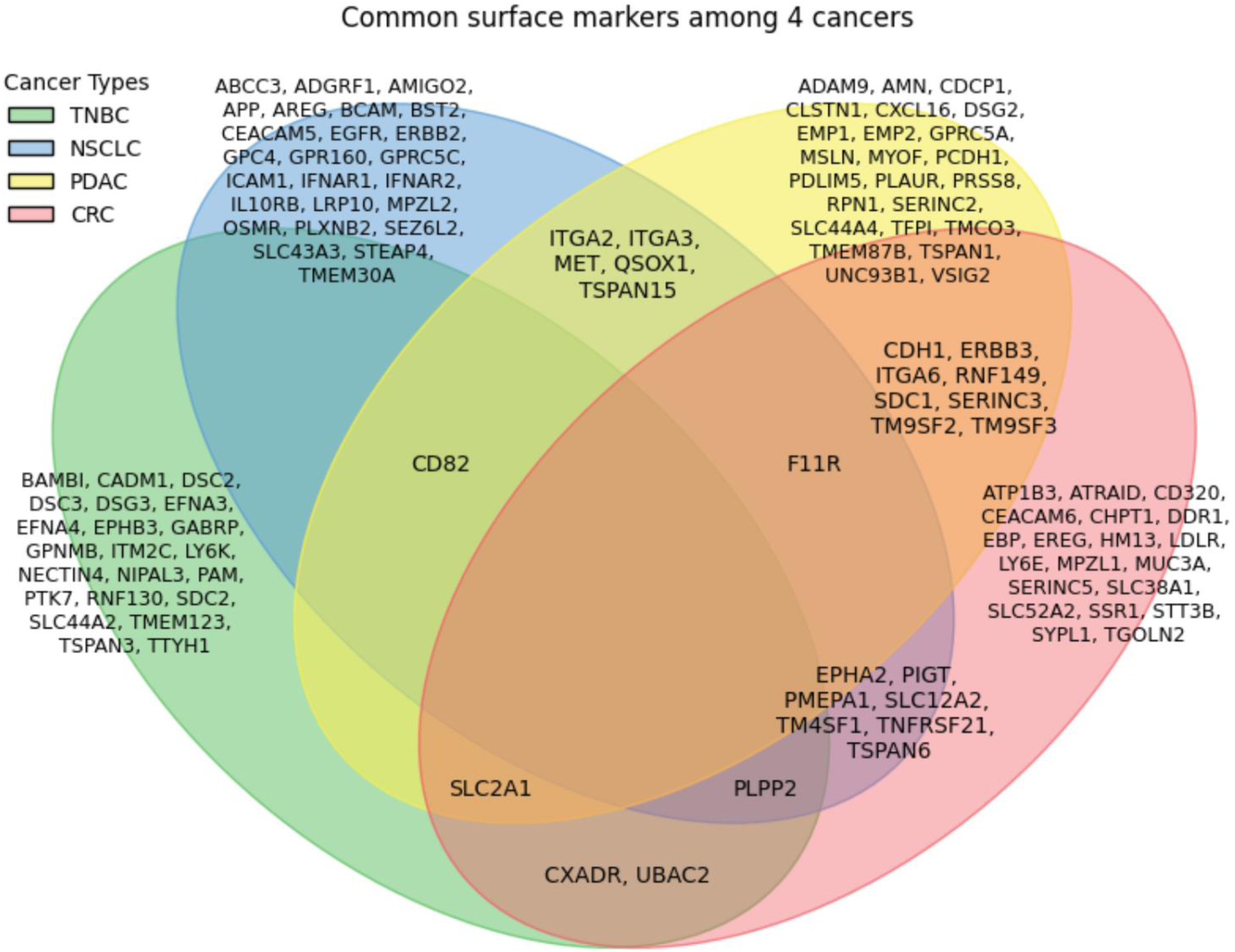
Venn diagram showing the overlap of surface marker genes among four cancer types: triple-negative breast cancer (TNBC), non-small cell lung cancer (NSCLC), pancreatic ductal adenocarcinoma (PDAC), and colorectal cancer (CRC). For each cancer type, the top 40 surface markers were selected based on marker scores (see Assessment of marker scores in the supplementary file 1), and their intersections were used to identify genes shared across multiple cancers. Although no markers were found to be common across all four cancer types, several notable markers were shared among three: *CD82* (TNBC, PDAC, NSCLC), *F11R* (PDAC, CRC, NSCLC), *SLC2A1*, *TM9SF2* (TNBC, PDAC, CRC), *CXADR*, and *PLPP2* (TNBC, NSCLC, CRC). Other markers were either uniquely associated with individual cancer types or shared between two.

Notably, several genes were shared among three cancer types: *CD82* (TNBC, PDAC, NSCLC), *F11R* (PDAC, CRC, NSCLC), *SLC2A1* (TNBC, PDAC, CRC), and *PLPP2* (TNBC, NSCLC, CRC). *CD82* has traditionally been described as a metastasis suppressor^62^, yet in our analysis it was consistently upregulated in tumor epithelial cells across three cancers, suggesting a context-dependent or state-specific function possibly unique to aneuploid subpopulations. *F11R* (JAM-A) is implicated in cancer cell adhesion and migration^47,63^ and has been associated with therapeutic resistance in several tumor types. *SLC2A1* (GLUT1) is a major glucose transporter frequently overexpressed in cancers and associated with metabolic reprogramming; it has been proposed as a diagnostic and therapeutic target in tumors including CRC and NSCLC^64,65^. *PLPP2*, a phospholipid phosphatase family member, has been shown to promote tumor proliferation and EMT-related pathways in lung adenocarcinoma, breast cancer, and PDAC, and is emerging as a potential lipid-signaling target in cancer therapy^66^.

These results reinforce the biological relevance of InferPloidy-driven target discovery. Not only were we able to recover well-established therapeutic targets, but we also identified potentially novel markers recurrently expressed across tumors. This highlights the power of integrating CNV-based tumor cell classification with expression-based prioritization for systematic surfaceome profiling. Together, our findings demonstrate that InferPloidy enables reliable identification of tumor-specific surface markers with high translational potential, supporting its application in therapeutic target discovery pipelines for precision oncology.

As a further investigation of inter-tumoral heterogeneity, we searched for cytogenetic bands with putative copy number amplifications (**Figs. 7 and 8**) using the method described at the end of the Implementation section (see also **Fig. 9**). Notably, the identified bands contained several well-known oncogenes, including *SOX2*^67^, *EIF3E*, *INTS8*^68^, *GSDMD*^69^, *NFASC*^70^, *ECT2*^67^, *EGFR*, and *ERBB2*, with the first four showing copy number amplification in three or more cancer types at an occurrence frequency exceeding 50% (**Fig. 8**). *ERBB2* (HER2), a therapeutic target in HER2^+^ breast cancer, was copy number amplified in all four HER2^+^ patients (**Fig. 8a**). Similarly, *EGFR*, a well-established target in NSCLC, exhibited copy number amplification in 8 of 9 NSCLC patients, albeit with variable amplitudes (**Fig. 8b**), and was also identified in PDAC and CRC with a frequency above 50 and 80%, respectively (**Fig. 8c** and **d**). *KRAS* also showed CNV gain in 5 out of 11 PDAC patients (**Fig. 8c**). Although these CNV profiles are inferred from gene expression data, this analysis demonstrates that InferPloidy enables cancer type- and subtype-specific CNV profiling by precisely distinguishing aneuploid from diploid cells.

**Figure 7.**
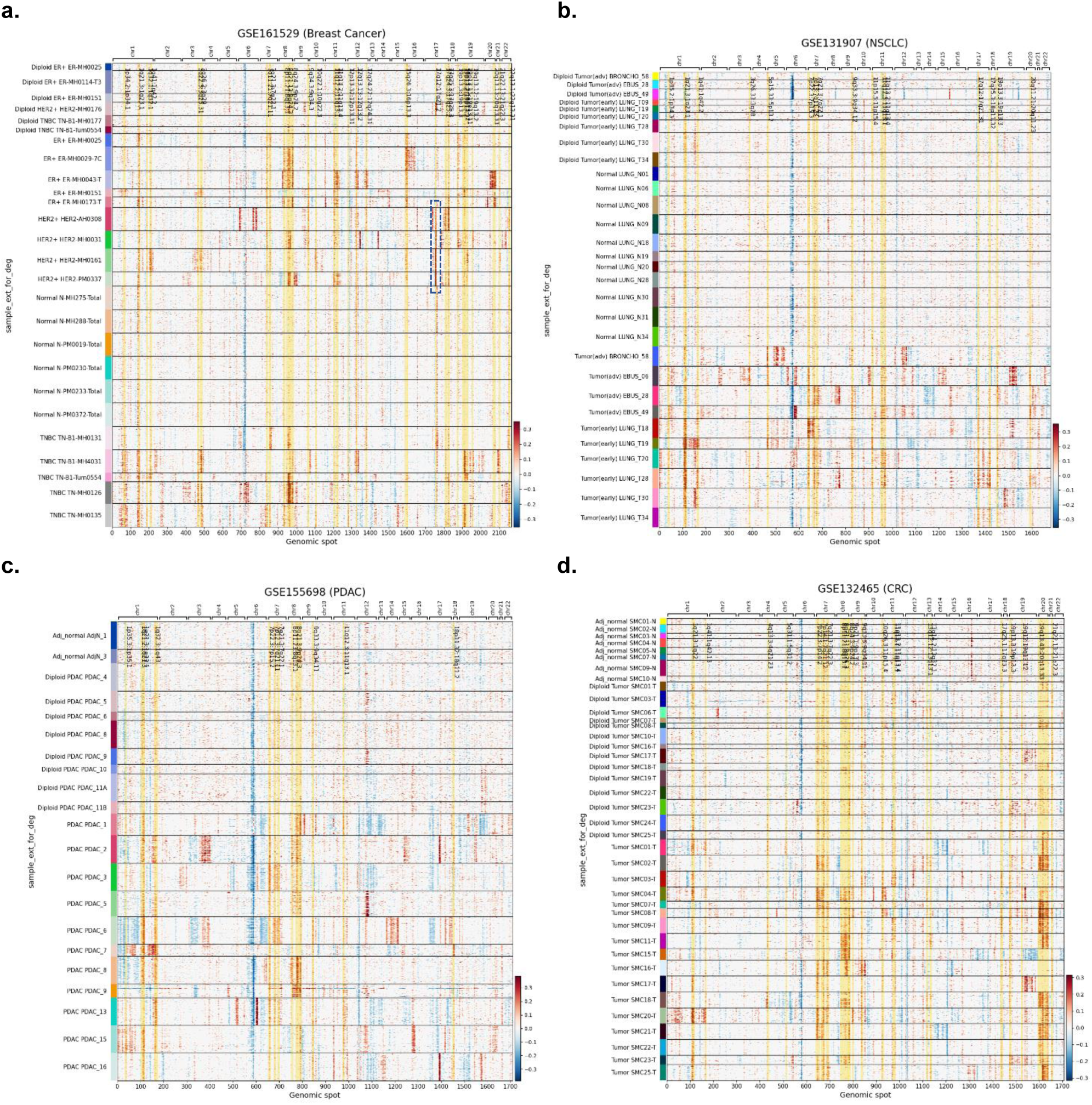
A comparison of CNV patterns in tumor-originating cells across four cancer types: (a) breast cancer, (b) non-small cell lung cancer (NSCLC), (c) pancreatic ductal adenocarcinoma (PDAC), and (d) colorectal cancer (CRC). The vertical axis represents cells grouped by sample (patient), and the horizontal axis represents genomic spot at which CNV was evaluated. Yellow-shaded regions indicate cytogenetic bands with significant CNV gain (≥4 samples) within a cancer type or subtype. In the four HER2^+^ breast cancer samples in (a), a pronounced gain is observed at the 1755^th^ genomic spot on chromosome 17, where the *ERBB2* gene is located (see also Fig. 8a).

**Figure 8.**
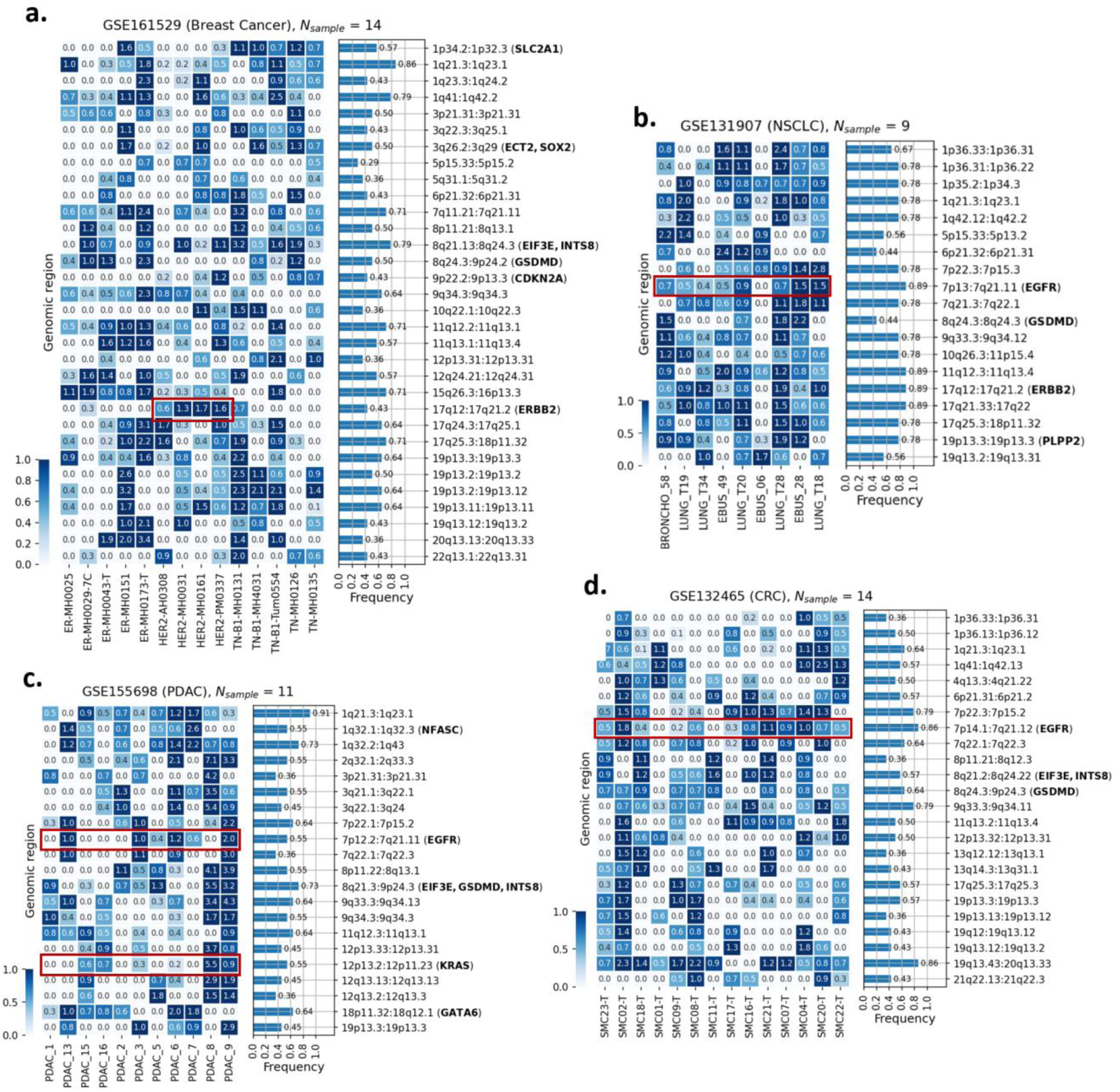
Summary of cytogenetic bands showing copy number gains in ≥4 samples for each cancer type/subtype: (a) breast cancer, (b) non-small cell lung cancer (NSCLC), (c) pancreatic ductal adenocarcinoma (PDAC), and (d) colorectal cancer (CRC). The heatmap depicts the average CNV gain score, *z*(*s*, *p*) (see Implementation) for each band, with darker colors indicating higher score. Bands containing well-known oncogenes, such as *EGFR* and *ERBB2*, are labeled in bold. The number of samples displayed is smaller than in the original datasets because some samples contained only diploid cells or had too few aneuploid cells for reliable identification of CNV gain region.

**Figure 9.**
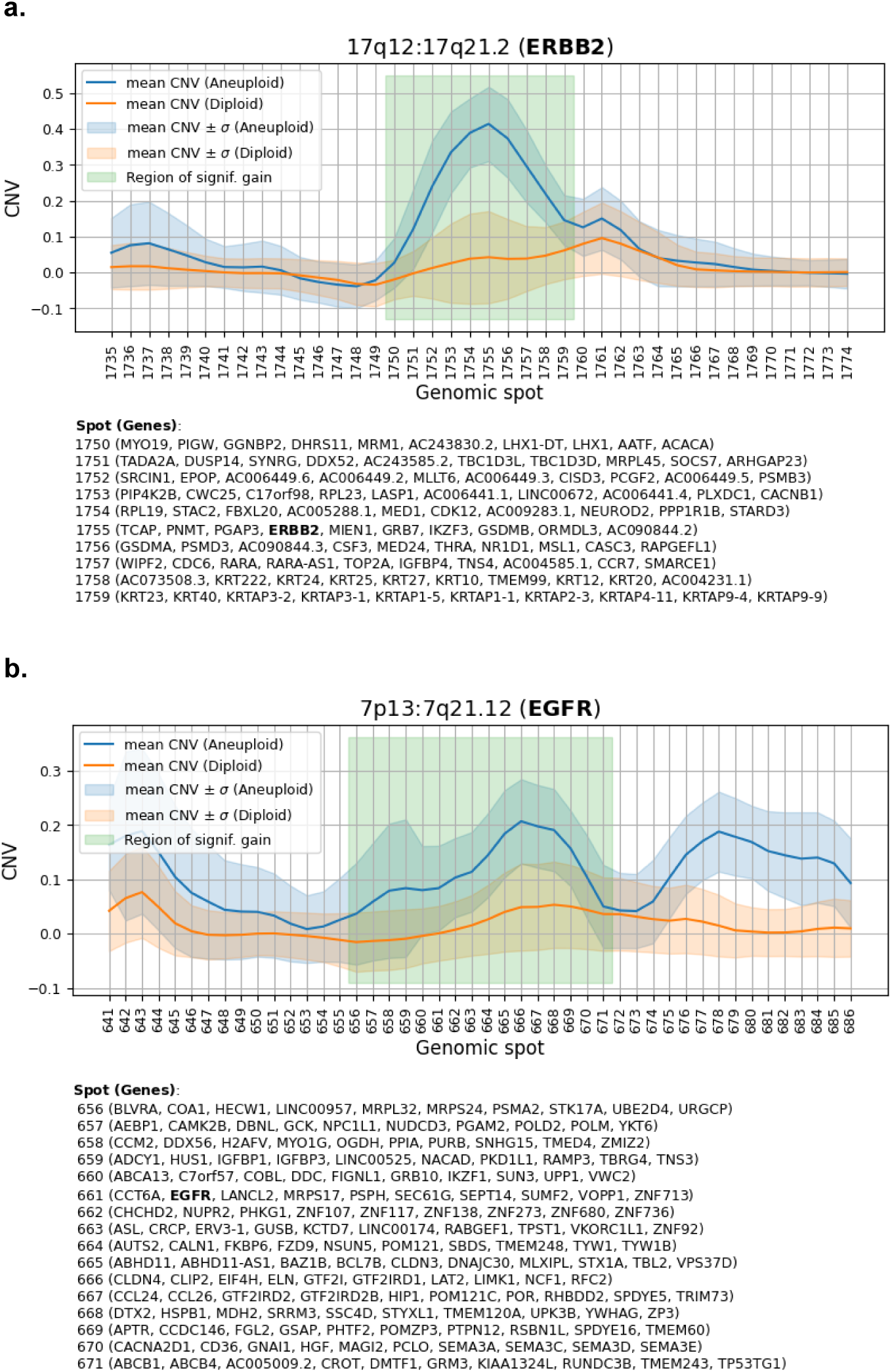
A brief description of the method to detect the region with significant copy number gain: (a) HER2^+^ Breast cancer and (b) non-small cell lung cancer (NSCLC). The horizontal axis indicates genomic spots; genes located in each spot are shown at the bottom.

## Conclusion

In this study, we present InferPloidy, a fast and accurate ploidy inference tool designed to complement the widely used CNV estimation platform, InferCNV. By combining a graph-based reference cell identification strategy with iterative Gaussian mixture modeling, InferPloidy achieved consistently higher classification accuracy than existing approaches such as CopyKat and SCEVAN, while reducing computational time by nearly two orders of magnitude (**Fig. 2–4**). This improvement removes a key bottleneck in CNV-guided tumor cell identification, enabling efficient analysis of large-scale, multi-sample single-cell RNA-seq datasets and facilitating integration into high-throughput analytical pipelines.

In datasets such as GSE161529, where certain samples contained relatively few normal reference cells, InferPloidy still maintained high classification accuracy by leveraging cluster structure and reachability within the CNV-derived expression space. While other tools such as CopyKat and SCEVAN can also operate without explicitly defined references, our comparisons indicate that InferPloidy provided more consistent results across varying sample compositions, underscoring its robustness in practical single-cell analysis settings. Although the current version does not aim to resolve clonal substructures, its scalability and reliability make it well suited for studies of tumor heterogeneity, cancer subtyping, and translational target discovery.

From a biological perspective, accurate separation of aneuploid tumor cells from diploid normal-like cells enhances the contrast between tumor-specific and background expression in downstream analyses, enabling cleaner differential expression results. Applied to four major cancer types—breast cancer, NSCLC, PDAC, and CRC—InferPloidy recovered well-established therapeutic targets such as *ERBB2* (HER2⁺ breast cancer), *ESR1* (ER⁺ breast cancer), *EGFR*, and *MET* (NSCLC), consistent with established targeted therapies (**Fig. 5**). Furthermore, cross-cancer comparisons revealed recurrently upregulated surface markers including *CD82*, *F11R*, *SLC2A1*, and *PLPP2* (**Fig. 6**), which have documented preclinical or clinical relevance. Beyond individual marker recovery, InferPloidy also enabled cancer type- and subtype-specific CNV profiling, revealing recurrent amplifications of well-known oncogenes such as ERBB2 and EGFR, as well as others shared across multiple cancer types, thereby providing an additional layer of insight into inter-tumoral heterogeneity (**Fig. 7** and **8**).

Overall, these findings highlight the utility of combining CNV-based tumor cell classification with systematic surfaceome profiling for marker and therapeutic target identification. By recovering both established and potentially novel candidates, InferPloidy offers a scalable and biologically informed framework for precision oncology. Future extensions incorporating clonal architecture analysis and integration with spatial transcriptomics may further expand its capability to dissect tumor ecosystems and inform next-generation therapeutic strategies.

## Data and Code availability

All datasets used in this study (GSE176078, GSE103322, GSE72056, GSE161529, GSE131907, GSE155698, and GSE132465) are publicly available in the Gene Expression Omnibus (GEO). The InferPloidy Python package is available on PyPI (pypi.org), and the full source code with <u>example Jupyter notebook</u> is hosted at https://github.com/combio-dku/InferPloidy. The processed data, including all results used to generate the plots in this paper, have been deposited in Figshare (https://figshare.com/articles/dataset/29886995). For reproducibility, the git repository also includes <u>the jupyter notebook</u> that downloads the processed data and reproduces the figures (Figs. 2–9).

## Authors’ contribution

SY, SJ and JM designed the overall experiments and devised the key idea. SY developed InferPloidy. DH, WS and JC performed single-cell RNA-seq data analysis. SY and JM guided the analysis. JL and SJ gave biological/medical interpretation and feedback on the results. All authors wrote, read, and approved the final manuscript.

## Acknowledgements

This research was supported by Basic Science Research Program through the National Research Foundation of Korea (NRF) funded by the Ministry of Education, Science and Technology (RS-2025-16070308) and the MSIT (Ministry of Science and ICT), Korea, under the ITRC (Information Technology Research Center) support program (IITP-2024-RS-2024-00437102) supervised by the IITP (Institute for Information & Communications Technology Planning & Evaluation)

